# Beyond early development: observing zebrafish over 6 weeks with hybrid optical and optoacoustic imaging

**DOI:** 10.1101/586933

**Authors:** Paul Vetschera, Benno Koberstein-Schwarz, Tobias Schmitt-Manderbach, Christian Dietrich, Wibke Hellmich, Andrei Chekkoury, Panagiotis Symvoulidis, Josefine Reber, Gil Westmeyer, Hernán López-Schier, Murad Omar, Vasilis Ntziachristos

**Affiliations:** Institute of Biological and Medical Imaging (IBMI), Helmholtz Zentrum München, Neuherberg, Germany; Chair for Biological Imaging and Translatum, Technische Universität München, München, Germany; Carl Zeiss AG, Corporate Research and Technology, Jena, Germany; Institute of Developmental Genetics, Helmholtz Zentrum München, Neuherberg, Germany; Research Unit Sensory Biology and Organogenesis, Helmholtz Zentrum München, Neuherberg, Germany

**Author notes:** These authors contributed equally to this work (Paul Vetschera, Benno Koberstein-Schwarz). Correspondence should be addressed to V.N.

## Abstract

Zebrafish animal models have traditionally been used in developmental biology studies but have recently become promising models of cancer, tissue regeneration and metabolic disorders, as well as efficient platforms for functional genomics and phenotype-based drug discovery. Most studies of zebrafish have examined only the embryonic or larval stages of development, yet many questions in developmental biology and biomedicine require analysis of adults, when zebrafish are large and opaque. Conventional microscopy methods are highly sensitive to light scattering and therefore cannot be applied to zebrafish older than a few weeks. We describe a novel multi-modality system that can observe zebrafish from the larval stage to adulthood. Using a hybrid platform for concurrent selective plane illumination microscopy (SPIM) and optoacoustic mesoscopy we show continuous imaging of fish growth over 47 days of development at a similar object size-to-resolution ratio. Using multiple wavelength illumination over the visible and short-wavelength infrared regions, we reveal that the optoacoustic method can follow GFP-based contrast used in SPIM, enabling molecular imaging interrogation in adult fish. Moreover we optoacoustically reveal many other features of zebrafish based on optical contrast not present in SPIM, including contrast from endogenous blood, water and lipids. The hybrid method presented can extend optical imaging to adult zebrafish employed as model systems for studying long-term processes in development, cancer, diabetes and other disorders.

## Introduction

Zebrafish (*Danio rerio*) is a widely used organism to address questions in developmental biology because it can be bred in large numbers in a cost-and space-effective way, it can be analyzed in detail at early developmental stages using high-resolution optical techniques, and it can be genetically manipulated to explore diverse cellular processes. More recently, zebrafish have become models of cancer, tissue regeneration and metabolic disorders^1^. They can provide insights into tumor growth, angiogenesis and metastasis^2^. They may even support the development of personalized anti-cancer medicine: patient-derived xenografts can be grown much faster in zebrafish and with much less patient material than in mice^3^. They can be used in drug activity screens based on alterations in phenotype that can be sensitively detected, including changes in pain, tumor metastasis, vascular tone and gut motility^4^. Similarly, they can be used to examine toxic effects of potential drugs or environmental contaminants on specific organs, and to screen potential neuroactive compounds for their ability to modify behavior. For instance, zebrafish express ghrelin and leptin, and recapitulate key aspects of obesity and type 2 diabetes^5^. The availability of a high-quality zebrafish genome facilitates detailed genetic analysis and gene-specific modifications.

These exciting applications of zebrafish to health and disease have so far been limited to studies of the embryonic and larval stages, when the fish are small (thin) and transparent and therefore amenable to analysis with high-resolution optical techniques. While these studies have provided numerous insights in development and disease, they cannot provide detailed structural information about long-term disease progression or long-term toxicity of drugs or environmental contaminants. Disease onset and pathophysiology can vary with age, such as in metabolic disorders^5^. In addition, morphogenesis and tissue remodeling continue throughout adulthood. These considerations make studies in adult zebrafish as important as those in earlier stages^6^.

Imaging adult zebrafish is challenging because their size and thickness preclude the use of purely optical imaging methods, as light cannot penetrate into deep tissue due to scattering. Depending on the microscopic technique and the extent of scattering, light microscopy can penetrate to depths of 0.1 – 1 mm ^7^, while the body thickness of model organisms such as zebrafish ranges from less than 100 µm in the early larval stage to nearly 1 cm in adult specimens. Selective plane illumination microscopy (SPIM) has been proposed for imaging specimens up to the late larval stage. By illuminating tissue using a thin sheet of light passing through the specimen, SPIM offers a simple way to achieve fast, volumetric imaging of living organisms, as long as the sample is small and transparent. The comparably low laser energy used in SPIM together with the fast acquisition speed makes it well suited for *in vivo* imaging. Multiple studies^8^ have already shown the value of SPIM for imaging small model organisms, such as the *in vivo* SPIM imaging of the beating heart of a 5-day-old zebrafish.^9^ However, SPIM imaging is not feasible in older, larger zebrafish, since SPIM resolution rapidly degrades due to aberrations for specimens thicker than 200 µm. Moreover, zebrafish lose their transparency as they grow, primarily caused by light scattering. Larger samples can be chemically treated to render them optically clear for SPIM imaging, but such treatment is toxic and does not allow *in vivo* measurements^10^. An engineered zebrafish strain called *crystal* is translucent as an adult and has proven useful in cancer studies^11^, but the genetic alterations in this strain are not entirely understood and may interfere with interpretation of results^2^.

Optoacoustic imaging has been considered for optical imaging beyond the depths reached by microscopy^7^. The method can image through opaque tissues and form images that are insensitive to scattering. In contrast to microscopy methods, which use a focused light beam vulnerable to scattering, the resolution in optoacoustic tomography images depends on the ultrasound frequencies detected ^7^,^12^,^13^. By utilizing ultra-wide bandwidth detection spanning at least 10-200 Mhz we have recently demonstrated the collection of rich information contrast from tissues in the mesoscopic regime at resolutions of a few to tens of microns at depths of a few millimeters in tissue^12^–^20^. For example, a 1-week-old zebrafish has been imaged with raster-scan optoacoustic mesoscopy (RSOM) at a lateral resolution of 18 µm in the 20-180 MHz frequency range^15^, while an axial resolution as high as 4 microns has been observed at the higher end of the frequency spectrum. The technique has also been used to image *Drosophila* in larval and adult stages^12,21^. One of the advantages of RSOM is that multiple excitation wavelengths can be used in order to image various sources of contrast, such as blood, melanin^14,17^, lipids and water^22^–^24^, yet this multi-wavelength feature has rarely been used for imaging small organisms^14,17^. Despite its advantages for zebrafish imaging at later stages of development, RSOM offers a one-sided view of the imaging subject, which reduces image quality as it cannot resolve objects that are parallel to the detection axis. In addition, the method typically collects data after illumination at only one or two wavelengths, limiting the amount of information that can be gained.

Here we aimed to combine, into a hybrid system, mesoscopy implemented in 360-degree multi-projection optoacoustic tomography with SPIM, which operates at a similar resolution. We hypothesized that the hybrid system could advance our capacity for longitudinal studies of small organisms, breaking through the fundamental scattering limit of purely optical methods. A question during such development was whether we could maintain imaging of the same contrast, i.e. fluorescent proteins, as we transitioned from SPIM to optoacoustic mesoscopy. Another question we investigated was what features could be resolved by optoacoustics at different illumination wavelengths, complementing features imaged by SPIM. We show that SPIM offers cellular resolution and forms images based on expression of green fluorescent protein (GFP) at early stages of development, while optoacoustic mesoscopy continues to detect GFP post embryogenesis, including when the organism is too thick or opaque to permit SPIM imaging. Moreover, we observe features of fish such as vascular contrast and water and lipid contrast. The integration of the two modalities into a single system allows the corresponding images to be accurately co-registered, and the 360-degree projections allow the acquisition of extensive optoacoustic data, leading to accurate images. Such imaging performance was achieved through the use of a novel sample holder and detector system design. The method proposed can become an invaluable tool in the application and study of zebrafish as an emerging tool for accelerating biological discovery and clinical translation.

## Results

The hybrid implementation was based on a common imaging chamber housing a SPIM and an optoacoustic mesoscopy system (**Fig. 1**). To accommodate both modalities and enable full 360° viewing for optoacoustic mesoscopy, a new SPIM design was introduced that enables multi-orientation scanning and illumination over the entire visible range from 420 to 700 nm. To integrate optoacoustic mesoscopy into the system, a homogeneous beam profile sized 10 × 10 mm^2^ was created in a second illumination path by using a diffusor and light-mixing rod. The sample is homogeneously illuminated from two sides and rotated over a full angle range of 360° during optoacoustic mesoscopy acquisition, while the linear transducer array is translated over the entire field of view (FOV) at each angle position. The central frequency of the linear array was chosen to be 24 MHz to provide similar resolution as SPIM. The cylindrically-focused 128 elements detect ultrasound signals from a distance of ∼ 8 mm from the sample center. As a result, the optoacoustic mesoscopy modality enables volumetric imaging with a FOV of 10 × 10 × 10 mm. An optical parametric oscillator (OPO) laser with a broad spectral range (420 to 2300 nm) is used to deliver nanosecond pulses, which stimulate acoustic emission from several absorbing tissue compounds, such as melanin, hemoglobin, water and lipid. During sample analysis with this hybrid system, typically SPIM images are first acquired, followed by optoacoustic mesoscopy scanning. Acoustic signals are volumetrically reconstructed using a filtered backprojection algorithm. Finally, SPIM and optoacoustic mesoscopy images can be co-registered based on the positions of the translation and rotation stages during data acquisition.

**Figure 1:**
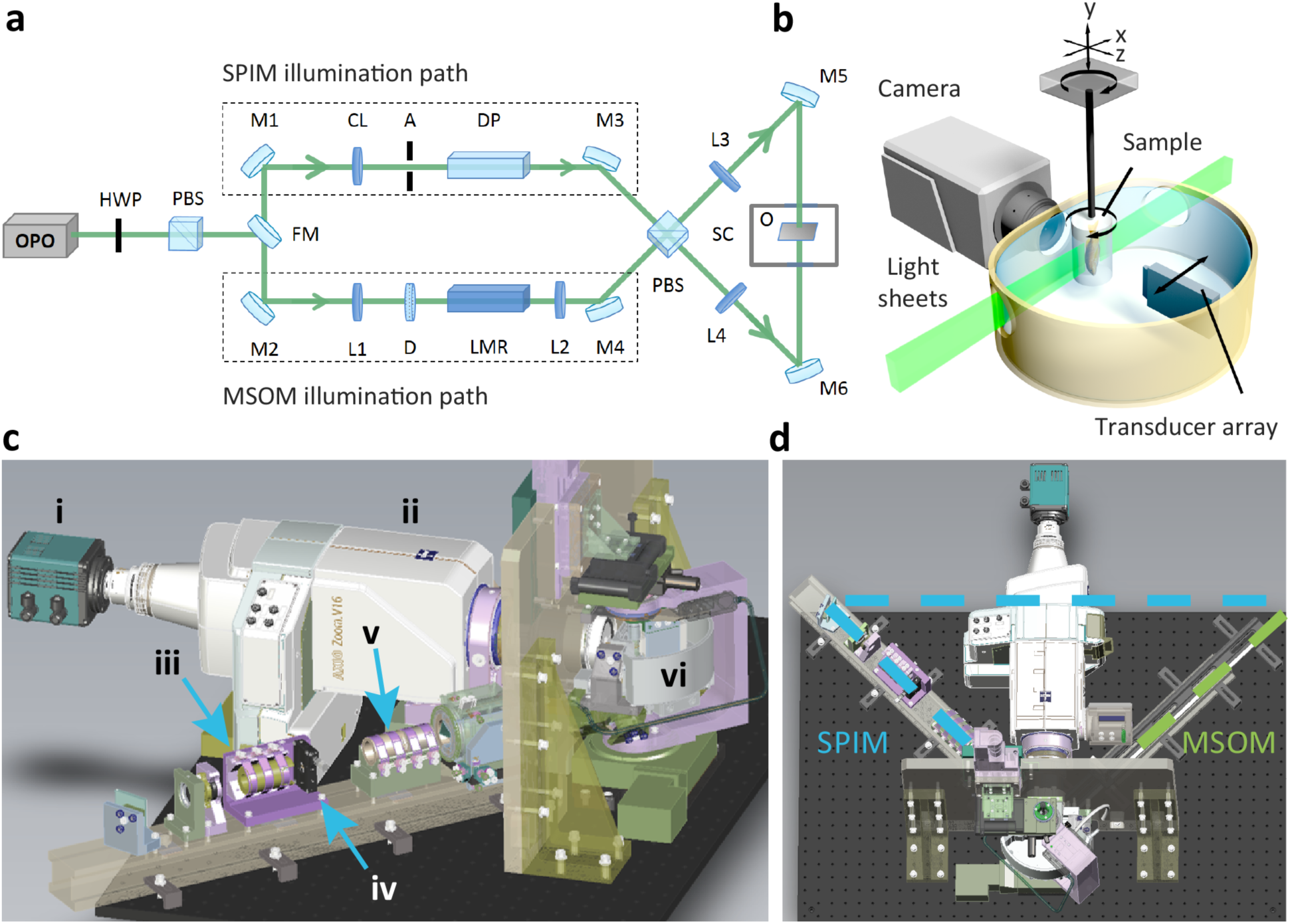
Schematic of the SPIM-optoacoustic mesoscopy set-up. **(a)** A pulsed optical parametric oscillator (OPO) laser illuminates first a half-wave plate (HWP) and a polarizing beam splitter (PBS) for power control and then a flip mirror (FM), which is used to choose between light sheet illumination for SPIM or broadfield illumination for optoacoustic imaging. The light sheet illumination involves a cylindrical lens (CL), aperture (A) and dove prism (DP); volumetric illumination for optoacoustic mesoscopy involves a lens (L1), diffusor and light mixing rod. The pulse energy is adjusted using a motorized half-wave plate (HWP) and a polarizing beam splitter (PBS); beam size is optimized using lenses L2-L4. A second PBS splits the beam 50:50 before illuminating the object (O) in the sample chamber (SC). **(b)** Enlarged view of the sample chamber. The sample is placed in an agar block, which is rotated around the indicated axis to provide multiple views and to allow optoacoustic tomography. In SPIM mode, the sample is linearly translated through the light sheets and subsequently rotated by 90° for multiple views. During optoacoustic acquisition, the sample is rotated 360° degrees and the transducer array is linearly translated in the x-direction at each angle position. **(c)** CAD-generated side view of the hybrid system, showing the camera (i), optical zoom (ii), cylindrical lens (iii), aperture (iv), diaphragm prism (v) and sample chamber (vi). **(d)** CAD-generated top view of the hybrid system. The dashed lines show the two beam paths for SPIM (blue) and optoacoustic mesoscopy (green).

First we characterized the performance of the hybrid system using agar phantoms containing fluorescent and polystyrene beads. **Fig. 2a-b** show SPIM images of a 10-μm fluorescent bead, indicating resolution of approximately 35 μm along the z axis and 10 μm along the x and y axis **(Fig. 2e)**. The lateral resolution in the x-and y-directions is restricted by the optical zoom and the camera. For the largest possible FOV with the optical zoom, a lateral resolution of 6.15 μm can be achieved. **Fig. 2c-d** show optoacoustic mesoscopy images of a 20-μm polystyrene bead, indicating a resolution of 35 μm along the x and y axes, and 120 μm along the z axis **(Fig. 2e)**. **Fig. 2f** shows images of crossed sutures with a diameter of 20 μm, demonstrating the location-independent high resolution over the full FOV.

**Figure 2:**
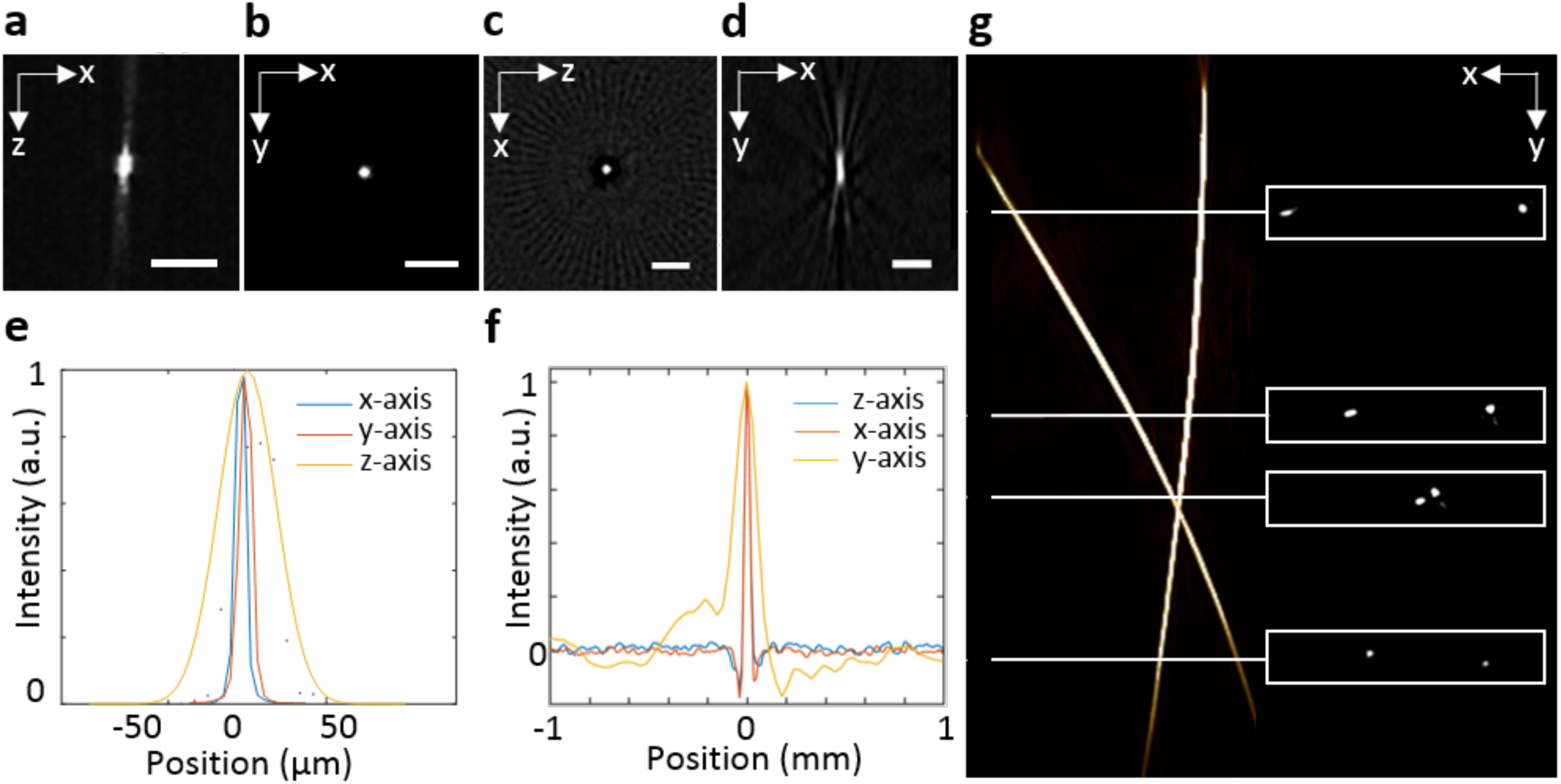
Characterization of the SPIM-optoacoustic mesoscopy hybrid system. **(a, b)** A 10-μm fluorescent bead recorded with SPIM. **(c, d)** A 20-μm polystyrene bead from two different perspectives as acquired with optoacoustic mesoscopy after reconstruction. **(e)** Profile plot for the 10-μm fluorescent bead measured with SPIM. **(f)** Profile plot for the 20-μm polysterene bead in the x-, y- and z-directions as measured with optoacoustic mesoscopy. The resolution of the optoacoustic mesoscopy modality was determined to be 35 × 35 × 120 μm based on full width at half-maximum in the profiles. **(g)** Full-view image of crossed 20-μm sutures acquired with the optoacoustic mesoscopy modality. The boxed insets show the cross-section from the top.

Next we used the SPIM-optoacoustic mesoscopy system to image Tg(pou4f3:GAP-GFP) transgenic zebrafish *(Danio rerio)* expressing GFP in hair cells and optic nerves from early development to adulthood. For these studies, we used a single excitation wavelength of 488 nm for both modalities. This wavelength allowed us to capture GFP fluorescence using SPIM as well as contrast from melanin and blood using optoacoustic mesoscopy. **Fig. 3a-h** shows SPIM- optoacoustic mesoscopy imaging of the GFP-expressing zebrafish from the larval stage (10 days) to adult stage (47 days). While abundant GFP signal with the expected distribution is visible in SPIM images of larvae, the signal is much weaker and less resolved in older animals. In contrast, the red optoacoustic signal from melanin and hemoglobin remains strong at all ages, and overall morphology as well as internal structures are visible. **Fig. 3k** shows the variance of GFP fluorescence as a function of imaging depth, which can be regarded as a simple and noise robust sharpness metric. The variance will increase for sharper images with higher intensity variations compared to a blurred image and can therefore be used to measure the amount of scattering in the sample. The plot shows that SPIM image sharpness rapidly decreases deeper than approximately 180 μm due to scattering. In contrast, the optoacoustic signal from the vessels identified in the zebrafish remain strong throughout the depth range scanned. As a result, the edges of blood vessels remain sharp even at depths of 400 μm (**Fig. 3l**). The measured full width at half-maximum of the depicted blood vessels was 35 µm in both cases, which is at the resolution limit of the optoacoustic mesoscopy system. These results highlight the complementary nature of the information obtained from SPIM and optoacoustic mesoscopy at a single wavelength.

**Figure 3:**
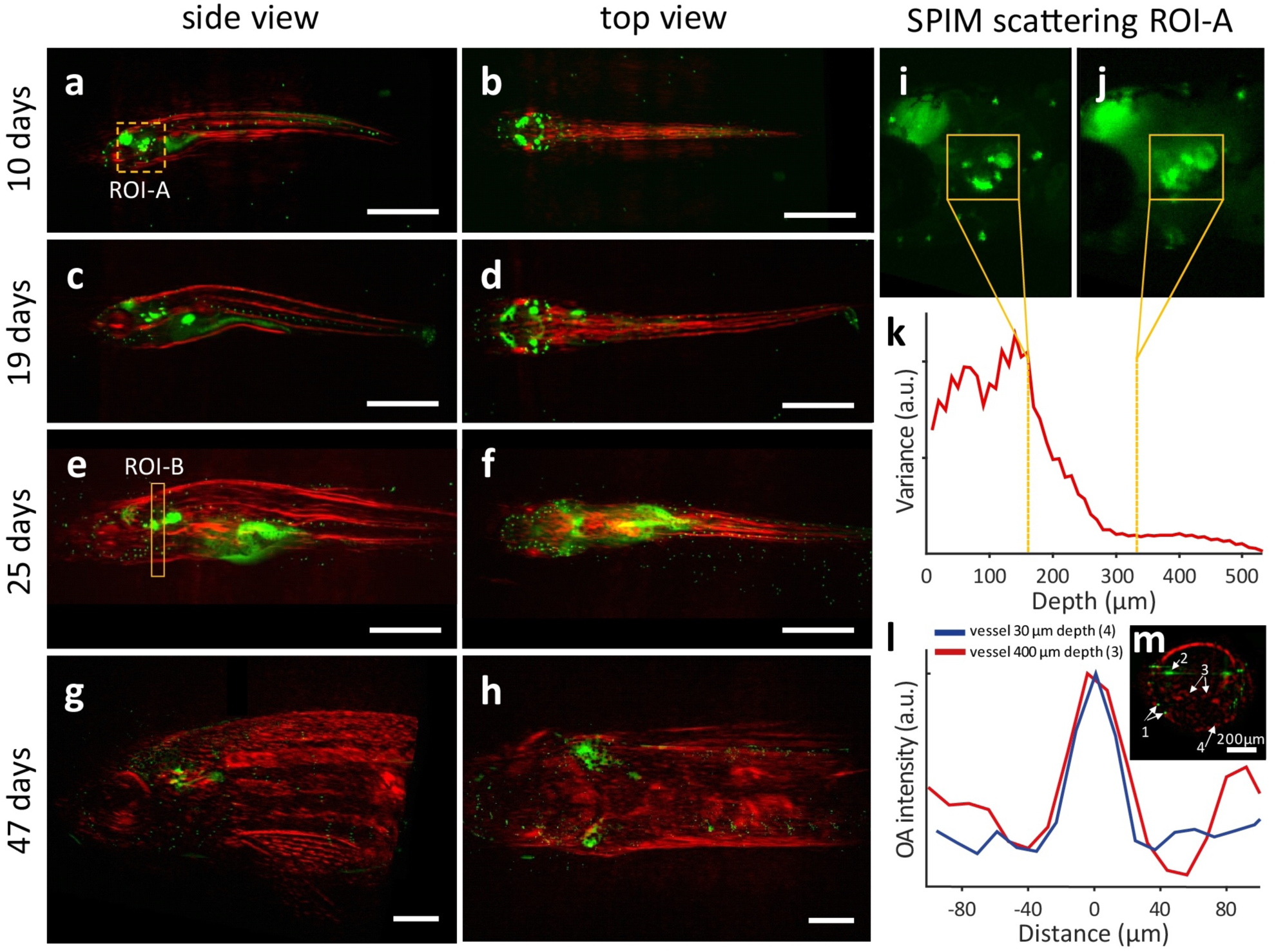
Hybrid SPIM-optoacoustic mesoscopy imaging of zebrafish development at a single wavelength. Transgenic zebrafish *(Brn3c:GFP)* expressing GFP in hair cells were imaged at 10, 19, 25, and 47 days after fertilization. GFP fluorescence (green) was imaged using SPIM, while the optoacoustic signal from melanin and hemoglobin (red) was reconstructed using optoacoustic mesoscopy. In both cases, a single excitation wavelength of 488 nm was used. **(a-h)** Side and top views of zebrafish at different developmental stages. A different fish was analyzed at each time point. **(i-j)** Two enlarged views of ROI-A in panel (a), with panel (i) lying at a depth of 160 µm (closer to the objective) and panel (j) lying at a depth of 340 µm. **(k)** Variance of fluorescence intensity as a measure of image sharpness. **(l)** Intensity profile of vessels imaged using optoacoustic mesoscopy, one at a depth of 20-30 µm below the surface [labeled “4” in panel (m)] and the other at a depth of 400 µm [labeled “3” in panel (m)]. **(m)** Cross-sectional view of ROI-B depicted in side view in panel (e). Numbers annotate anatomical features: 1, hair cells; 2, ear; 3 and 4, vessels. Scale bars in panels (a-h) indicate 1 mm.

Next we focused on the ability of optoacoustic mesoscopy to complement the fluorescence contrast of SPIM. The multiwavelength capability of optoacoustic mesoscopy allows imaging based on several endogenous sources of contrast, which means that anatomical as well as functional information can be gained non-invasively for early and late developmental stages. Moreover these images can be co-registered perfectly with SPIM images to provide additional, fluorescence-based contrast. These optoacoustic mesoscopy experiments were conducted at two wavelengths to provide contrast due to melanin and hemoglobin. **Fig. 4a** and **4b** depict side and top views, respectively, of zebrafish, while **Fig. 4c-e** show the images obtained after spectrally unmixing the reconstruction data to separate the contributions of melanin and hemoglobin. The unmixing revealed several anatomical features, including the hyomandibula, gill filaments, dorsal aorta and cardinal vein.

**Figure 4:**
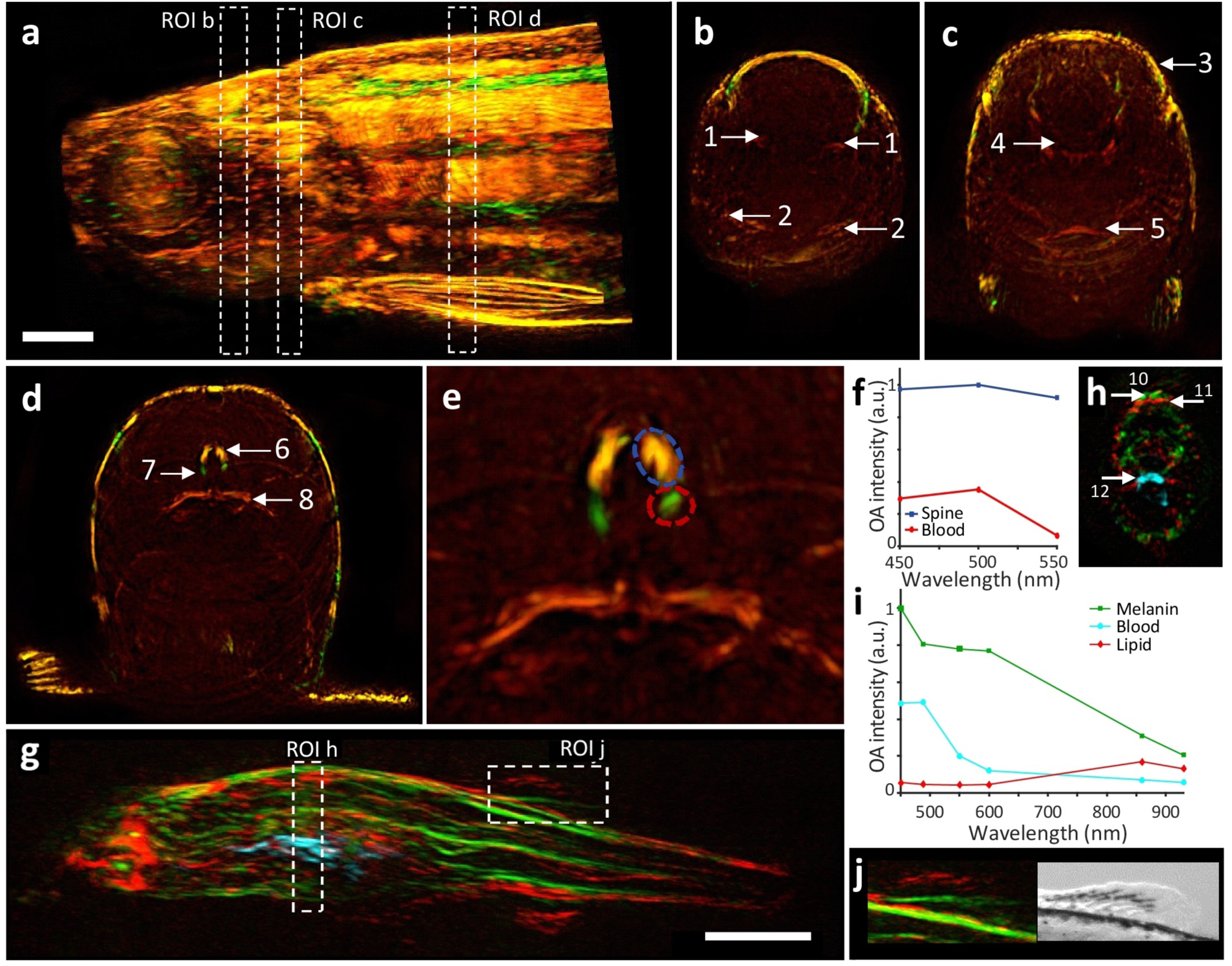
Optoacoustic mesoscopy of zebrafish at multiple wavelengths. **(a-e)** A wild-type zebrafish not expressing GFP was imaged at 2 months old after illumination at 450 nm (green) and 550 nm (red). Reconstructions at each of these wavelengths were superimposed, such that overlapping features appear yellow. **(a)** Sideview maximum intensity projection (MIP). **(b-d)** MIPs showing the three ROIs from panel (a). Numbers annotate anatomical features: 1, hyomandibula; 2, gill filaments; 3, semicircular canal; 4, medulla oblongata; 5, cardiac ventricle; 6, spinal cord; 7, cardinal vein; 8, swim bladder. **(e)** Enlarged view of the region in panel (d) around the spine. **(f)** Three-wavelength spectra of the spine and vessel marked in panel (e). **(g-j)** A transgenic zebrafish (*Brn3c:GFP*) at 25 days old was imaged after illumination at 450, 488, 550, 600, 860 and 930 nm. Acoustic data were linearly unmixed and reconstructed such that melanin contrast appears green; lipids, red; and blood vessels, blue. (g) Sideview MIP. (h) Cross-sectional MIP showing the melanin pigmented skin (10), subcutaneous lipid (11) and blood in the region of the heart (12). **(i)** Unmixed spectra of melanin, blood and lipid. **(j)** Enlarged view of the upper fin from panel (g). *Inset*, a photograph of the same area. Scale bars in panels (a) and (g) indicate 1 mm.

To image even more sources of contrast, we expanded the excitation range of our imaging system to the near-infrared region (700-930 nm) and applied a linear multispectral unmixing algorithm using the intrinsic optoacoustic spectra of the sample (**Fig. 4f**). The resulting images in **Fig. 4g-h** reveal structures different from those observed with wavelengths in the visible region of the spectrum, including blood-containing cardiac structures deep within the fish **(Fig. 4g** and **4h**). We attribute the non-blood-containing structures to water and lipid, which absorb more strongly in the near-infrared region. Consistent with this interpretation, **Fig. 4j** shows that areas of optoacoustic melanin signal in the upper fin correspond to darkly pigmented areas in a photograph, while areas of putative optoacoustic lipid and water signal correspond to transparent areas. **Fig. 4i** shows the intrinsic spectra for melanin, blood and lipid structures, which provided the basis for linear unmixing.

Having demonstrated the ability of optoacoustic mesoscopy to complement the range of contrasts that can be imaged in juvenile and adult zebrafish, we wanted to examine whether optoacoustic mesoscopy could detect the same fluorophores as SPIM but in older zebrafish that cannot be imaged well with SPIM. To maximize the signal-to-noise ratio (SNR), we replaced the linear transducer array that we used for the abovementioned studies, which has small transducer elements of 0.07 × 3 mm^2^, with a large, spherically focused, single-element detector with a diameter of 10 mm. Tests with 50-μm polysterene beads embedded in an agar phantom indicated that this change in transducer increased SNR by ∼ 25 dB. To allow scanning of the sample over the full 360 degrees, in contrast to the limited angular range of previous optoacoustic mesoscopy set-ups^19^, we designed a sample holder that can be spirally rotated and translated in front of a stationary detector (**Fig. 5a**). A 45-day-old zebrafish (*cldnb:GFP, sqET4:GFP*) expressing GFP in the hair cells and the lateral line was imaged at 488 and 550 nm (**Fig. 5b**). The differential optoacoustic image in **Fig. 5e** was obtained by subtracting signals at 550 nm from signals at 488 nm, and it clearly reveals GFP expressed in the brain and lateral line in three dimensions throughout the sample. In contrast, **Fig. 5f** shows the average intensity projection of the acquired SPIM image stack for the same fish, revealing the comparatively diffuse GFP signal that overlaps with the signals observed by optoacoustic mesoscopy but with less resolution because SPIM is sensitive to scattered light. These results demonstrate that with an appropriately sensitive detector, optoacoustic mesoscopy can volumetrically image the contrast of genetically encoded fluorescent proteins in living model organisms with good resolution at developmental stages when SPIM functions poorly. **Fig. 5c** depicts the cross-section of the optoacoustic reconstruction marked in **Fig. 5b**, showcasing the depth-imaging abilities of the adapted system. **Fig. 5d** shows the absorption and optoacoustic spectra of GFP, which explain the strong difference in optoacoustic signal generation between 488 and 550 nm.

**Figure 5:**
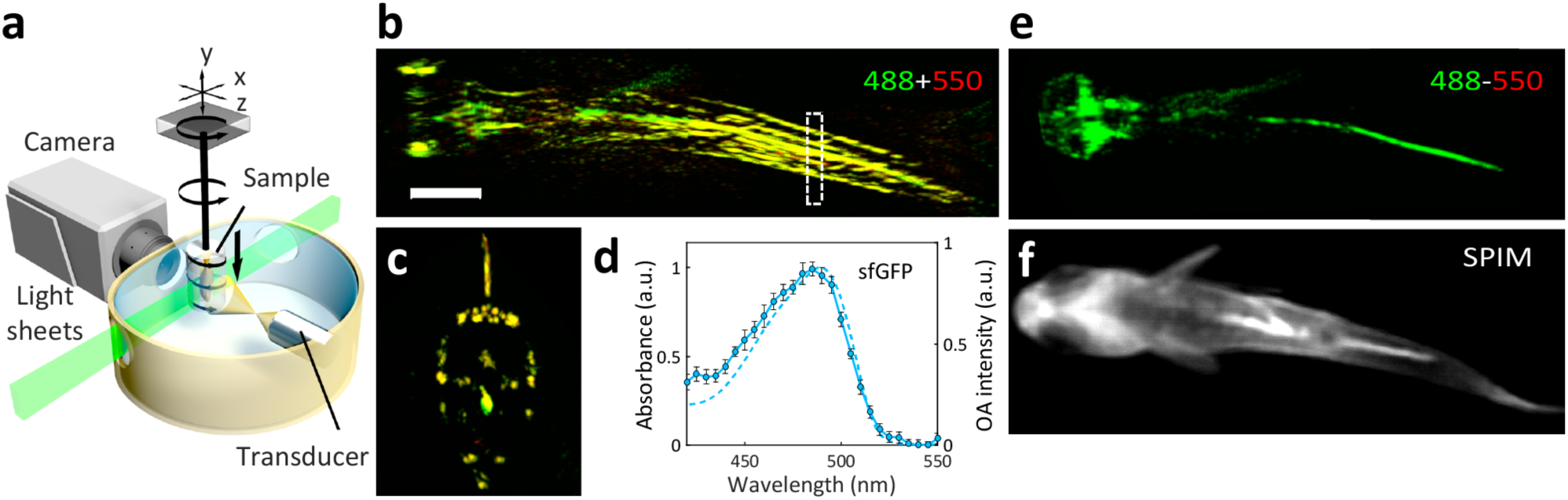
Spiral multispectral optoacoustic mesoscopy (SpiMSOM). **(a)** Enlarged view of the adapted sample chamber for the single-element transducer. During the optoacoustic mesoscopy acquisition, the sample is rotated 360° degrees and vertically translated downwards. Due to the broad numerical aperture of the transducer, the detector is able to acquire ultrasound signals from the complete sample width without any translation. **(b)** MIP of the top view of the superimposed optoacoustic reconstructions at 488 nm (green) and 550 nm (red) of a 45-day-old transgenic zebrafish (*cldnb:GFP, SqET4 GFP*). **(c)** Cross-sectional MIP of a region below the fin. **(d)** Absorbance and optoacoustic extinction spectrum of superfolder-GFP (see Methods for details). **(e)** Difference image (488 nm - 550 nm) of the optoacoustic reconstructions superimposed in panel (b). **(f)** Average intensity projection of the SPIM image stack of the same fish.

## Discussion

Here we describe a novel SPIM-optoacoustic mesoscopy imaging system that significantly improves on the imaging depths that can be achieved with conventional fluorescence microscopy, allowing the study of model animal development from early stages through adulthood. As we demonstrate here, the hybrid system can detect fluorescence contrast through SPIM and optoacoustic contrast of melanin, blood, lipids, fluorescent proteins and water. This allows multi-scale morphological and functional imaging throughout the zebrafish lifespan, positioning the hybrid system as a useful next-generation tool for developmental, cancer and neurobiological studies spanning all the life stages.

Among the hallmarks of cancer^25,26^ are angiogenesis, metastasis and metabolic reprogramming, and all these processes can be examined with optoacoustics^24^. One of the drivers of many cancer hallmarks is hypoxia within tumors, and optoacoustics can measure intra-tumor oxygenation with high spatiotemporal resolution^27^, even in response to perturbations^28^. Focal hypoxia in solid tumors may help drive tumor progression and therapy resistance^29^, so quantitating hypoxia with our hybrid system may allow *in vivo* screening of emerging anti-cancer therapies aimed at inhibiting tumor angiogenesis^30^. Indeed, our system might be well suited to exploring how hypoxia can drive angiogenesis and subsequently metastasis^31^. Our system is well-positioned to exploit transgenic and genome-modifying tools such as CRISPR/Cas^32^, for precise analysis of genes and proteins in healthy and diseased fish from embryonic stages through adulthood. Optoacoustics can also track rapid processes through detection of transgenically expressed reporter proteins, such as the calcium reporter GCaMP^33,34^.

Here, we develop an optoacoustic mesoscopy system in which the sample is rotated while the detector remains stationary. This provides better resolution and overall image quality than the previous optoacoustic mesoscopy system^19^, and allows deeper imaging within samples than a previous hybrid SPIM-optoacoustics system^35^. Our study is the first report of optoacoustic mesoscopy to examine a model organism at different stages of development. The unusual ability of optoacoustic mesoscopy to detect a range of developmentally relevant contrasts including, water, lipid and blood relies on excitation with a broad spectral range, in this case from 450 to 930 nm. This offers interesting potential for monitoring developmental changes e.g. in angiogenesis^36^, tissue regeneration^37^ cancer studies in zebrafish^3^. Our demonstration that optoacoustic mesoscopy can detect fluorescent proteins such as GFP even deep within older zebrafish means that this modality may be compatible with the myriad fluorescent proteins and transgenic expression systems that have been developed for optical microscopy studies^38^ but are limited to earlier developmental stages when the organisms remain transparent and thin. Our set-up may allow these tools to be extended to *in vivo* studies of adult stages. At the same time, the range of agents compatible with optoacoustic mesoscopy continues to grow^39^–^41^, opening up additional possibilities for *in vivo* imaging using our set-up.

Our novel hybrid system allows imaging of fluorescence and optoacoustic signal to visualize components and processes at the tissue and organismal levels. In addition, the multispectral capability of the system allows the distribution of numerous target molecules or tissues to be imaged simultaneously, either in a label-free way or following expression or injection of exogenous contrast agents. Our set-up is compatible with other model organisms such as *Xenopus* and axolotls, and it may be particularly useful for organisms that go from transparent to opaque during development and so cannot be analyzed at later developmental stages *in vivo* using conventional optical microscopy.

## Methods

### Experimental set-up

To provide the light sheet illumination for SPIM and homogenous illumination for optoacoustic imaging, a fast tunable nanosecond-laser was used (7 ns, 19 mJ at 450 nm, tunable from 420-700 nm, Spitlight-DPSS 250 ZHG-OPO, Innolas, Germany). The laser beam was aligned to the illumination paths for SPIM and optoacoustic mesoscopy (**Fig. 1**). A half-wave plate (HWP) and a polarizing beam splitter (PBS) were used to reduce the power to 25% for SPIM illumination. The light sheet for SPIM imaging was obtained using a cylindrical lens (CL) and an adjustable aperture (A), and consequently rotated by 90° using a dove prism (DP). The light sheet was divided at a second PBS to obtain double-sided illumination. With a flip mirror (FM) behind the first polarizing beam splitter, the beam was reflected to create a squared, homogeneous beam profile for optoacoustic imaging. Volumetric illumination was achieved by focusing the laser pulse on a diffusor (D), which was placed in front of a light mixing rod (LMR). The lens pairs L2-L3 and L2-L4 enlarged the squared beam profile to 10 × 10 mm^2^ in the sample chamber, which was again divided by the PBS for double-sided illumination. The maximum energy per pulse reaching the sample was measured as 3.5 mJ for both illumination paths.

### Hybrid SPIM and optoacoustic mesoscopy

To examine the combination of SPIM and optoacoustic mesoscopy for visualizing zebrafish development, *Brn3c:GFP* zebrafish of different ages were imaged *ex vivo* using SPIM to collect fluorescence signal from GFP, and also using single-wavelength optoacoustic mesoscopy at 488 nm to generate optoacoustic signal from melanin. Fish were scanned through the perpendicular light sheet on a translational stage (M-112.1DG; PI Micro, Germany). Fluorescence acquisition was performed with a scientific complementary metal oxide semiconductor (sCMOS) camera (pco.edge, PCO, Germany). Subsequently, the sample was rotated by 90°, then scanned again for multiview acquisition. Data were fused in Amira™ (Amira 6.2, Thermo Fisher Scientific, USA). Using fluorescent beads, we determined the minimal axial SPIM resolution to be approximately 35 μm. The lateral resolution in the xy-plane, determined by the optical zoom and the camera, was 6.15 μm.

Ultrasound detection of the samples in **Figs. 3** and **4** was performed using a cylindrically focused, wide-bandwidth ultrasonic transducer array (24 MHz center frequency, 55% bandwidth at −6 dB, 128 elements, Vermon, France). The translation-rotation scanning geometry was based on the concept presented by Gateau et al. ^18^. In contrast to the system described by Gateau et al.^18^, here the sample was rotated over 360° using a rotational stage (RS-40, PI Micro, Germany) and the ultrasound linear array was translated at each angular position with a linear stage (M-111.1DG, PI Micro, Germany). The sample was positioned using three linear stages in the x-, y-, and z-directions (M-112.1DG, M-403.2DG, and M-112.1DG, PI Micro, Germany). The system includes an additional rotational stage (M-061.PD, PI Micro, Germany) for rotating the array, which allows the detector to be moved so that the system can exploit the whole scanning range while running in SPIM mode.

To increase sensitivity for the optoacoustic imaging experiment in **Fig. 5**, the linear array was replaced with a spherically focused, single-element ultrasonic transducer (20 MHz center frequency, 77% bandwidth at −6 dB, Imasonic, France). In the novel spiral scanning geometry with the single-element detector, the sample is rotated continuously at the maximum speed of the rotation stage, while being slowly and simultaneously translated in the y-direction. The high numerical aperture of the detector enables detection of ultrasonic signals across the lateral width of the sample without any translation of the detector.

Following the optoacoustic scan, the raw signals were filtered using a third-order Butterworth filter, then reconstructed using a weighted backprojection algorithm^42^. For data used for linear unmixing, a Hilbert transform was applied to the data before the reconstruction in order to remove negative values and improve data quality for unmixing^18^. The weighted backprojection was performed on cuboid voxels with a size of 12 × 12 × 35 μm^3^ for array scans and 8 × 8 × 16 μm^3^ for single-element scans, which exhibit a lower number of projections. The 3D image stacks were obtained using maximum intensity projections (MIP) and visualized with Amira™. Characterization with polymer beads of 10-20 μm revealed a resolution of 35 × 35 × 120 μm in the x-, y-, and z-directions for the array and 35 × 35 × 100 μm for the single-element detector. These values are consistent with the specifications of our transducers.

### Multispectral optoacoustic mesoscopy (MSOM)

To examine the ability of our hybrid SPIM-optoacoustic mesoscopy system to provide optoacoustic information simultaneously from multiple components of tissue, we imaged wild-type adult zebrafish (2 months old, ZDB-GENO-960809-7) *ex vivo* using optoacoustic mesoscopy as described above, except that we illuminated the animals using light at 430, 450, 488 and 550 nm to stimulate the different optoacoustic spectra of hemoglobin and melanin. The raw optoacoustic data were spectrally unmixed using a linear regression method^43^. Extinction spectra of melanin and blood were extracted from the reconstruction volume’s intensity profiles in regions corresponding to the four wavelengths measured. When visualizing fluorescent proteins such as GFP, we overlaid image reconstructions at 488 and 550 nm **(Fig. 5b)** and subtracted the two reconstructions (488-500 nm; **Fig. 5e**). The optoacoustic spectrum shown in **Fig. 5d** was measured with the same laser but a different system described elsewhere ^44^.

## Acknowledgments

The authors wish to thank A. Chmyrov and A.C. Rodríguez for discussions on the manuscript. The authors thank L. Pola-Morell (Research Unit Sensory Biology and Organogenesis, Helmholtz Zentrum München, Neuherberg, Germany) for kindly providing mutant and wild-type zebrafish. The research leading to these results has received funding by the Bundesministerium für Bildung und Forschung (BMBF), Bonn, Germany (Project Tech2See, 13N12623, 13N12624). All authors declare that they have no competing interests.

